# Vertical niche definition of test-bearing protists (Rhizaria) into the twilight zone revealed by in situ imaging

**DOI:** 10.1101/573410

**Authors:** Tristan Biard, Mark D. Ohman

**Affiliations:** Scripps Institution of Oceanography, University of California, San Diego, La Jolla, CA 92093, USA; Laboratoire d’Océanologie et Géosciences, Université du Littoral Côte d’Opale, 28 avenue Foch, 62930 Wimereux, France

**Keywords:** Niche definition, In situ imaging, Rhizaria, Foraminifera, Phaeodaria, Collodaria, Acantharia, Protist, California Current Ecosystem, Vertical distributions, Mesopelagic.

## Abstract

The Rhizaria is a super-group of ameoboid protists with ubiquitous distributions, from the euphotic zone to the twilight zone and beyond. While rhizarians have been recently described as important contributors to both silica and carbon fluxes, we lack the most basic information about their ecological preferences. Here, using the in situ imaging (Underwater Vision Profiler 5), we characterize the vertical ecological niches of different test-bearing rhizarian taxa in the southern *California Current Ecosystem*. We define three vertical layers between 0-500 m occupied, respectively, by 1) surface dwelling and mostly symbiont-bearing rhizarians (Acantharia and Collodaria), 2) flux-feeding phaeodarians in the lower epipelagic (100-200 m), and 3) Foraminifera and Phaeodaria populations adjacent to the Oxygen Minimum Zone. We then use Generalized Additive Models to analyze the response of each rhizarian category to a suite of environmental variables. The models explain between 13 and 93% of the total variance observed for the different groups. While temperature and the depth of the deep chlorophyll maximum, appear as the main factors influencing populations in the upper 200 m, silicic acid concentration is the most important variable related to the abundance of mesopelagic phaeodarians. The relative importance of biotic interactions (e.g., predation, parasitism) is still to be considered, in order to fully incorporate the dynamics of test-bearing pelagic rhizarians in ecological and biogeochemical models.

## Introduction

Over the last two decades, interest has been elevated in the role of unicellular eukaryotes (i.e., protists) in marine global biogeochemical cycles (Worden et al. 2015). Among the large diversity of protists inhabiting the ocean (Massana 2015; Simpson et al. 2017), the Rhizaria is a diverse super-group of unicellular organisms, some of which have planktonic life styles (Burki and Keeling 2014). A typical feature of many marine rhizarians is the formation of a mineral test (or skeleton), whose chemical nature varies between taxa: Acantharia use strontium sulfate (SrSO_4_), Polycystine radiolarians and Phaeodaria secrete opaline silica (SiO_2_ nH_2_O), and Foraminifera form calcium carbonate (CaCO_3_) tests (Kimoto 2015; Nakamura and Suzuki 2015; Suzuki and Not 2015). With their mineralized skeletons eventually sinking upon death, test-bearing rhizarians actively contribute to fluxes of biogenic material into the deep ocean: Foraminifera are responsible for 32-80% of annual CaCO_3_ export to the deep ocean (Erez 2003), while Acantharia play a critical role in the ocean’s strontium budget (Bernstein et al. 1987) and Polycystines/Phaeodaria can act as major exporters of biogenic silica and particulate organic carbon, especially in oligotrophic regions (Lampitt et al. 2009; Guidi et al. 2016; Biard et al. 2018). Despite their active contribution to biogeochemical processes and substantial contribution to the marine carbon pool (Biard et al. 2016), there are still many critical gaps in our knowledge of test-bearing rhizarians, impeding full understanding of their ecological significance at local-to-global scales.

With fossil records as old as the lower Paleozoic (ca. 515 Ma), test-bearing rhizarians are among some of the oldest protist lineages found in modern oceans (Suzuki and Oba 2015). Through their long evolutionary history, they have adapted to a multitude of environmental conditions resulting in the ubiquity of contemporary rhizarians. They thrive from polar regions to warm equatorial waters, and from the sunlit surface to the dark meso- and bathypelagic ocean (Anderson 1983; Suzuki and Not 2015). A substantial fraction of test-bearing rhizarians inhabit the euphotic zone, hosting multiple photosymbionts from diverse eukaryotic and prokaryotic lineages (e.g., dinoflagellates, cyanobacteria) (Anderson 2012; Decelle et al. 2015). This mixotrophic behavior allow some of them to thrive in hostile environments such as oligotrophic gyres where food and nutrients are scarce (Taylor 1982; Biard et al. 2016). Deeper, in the dark ocean, are found deep-dwelling rhizarians (e.g., Phaeodaria) (Nakamura and Suzuki 2015), with various trophic modes. Some are carnivores feeding on mesopelagic plankton (e.g., Foraminifera) (Hull et al. 2011; Gaskell et al. in review) and others, flux feeders (e.g., phaeodarians), feeding on carbon rich particles raining from the euphotic layer (Gowing 1989). To date, most of the limited knowledge on rhizarian distribution patterns – both horizontal and vertical – is still inferred from surficial sediment samples, and little information is available on taxon-specific vertical niche distribution (Kling and Boltovskoy 1995). In addition, as with many protistan lineages (Caron et al. 2004), test-bearing rhizarians remain mostly uncultured (with the exception of some planktonic foraminifera; Kimoto 2015). Therefore, in situ experiments and observations are currently the only reliable alternative (apart from DNA surveys, which lack a true quantitative dimension) to improve knowledge on rhizarian ecology and vertical niche distribution.

Numerous in situ imaging ocean systems are available (Benfield et al. 2007), with different advantages and drawbacks, and sampling characteristics, ranging from towed systems (e.g., Video Plankton Recorder - Davis et al. 1992a; In Situ Ichthyoplankton Imaging System - Cowen and Guigand 2008), vertical profilers (e.g., Underwater Vision Profiler; Picheral et al. 2010), to fully autonomous underwater vehicles (e.g., *Zooglider*; Ohman et al. 2018). Despite different technical specifications, in situ imaging approaches all share the same potential to study delicate organisms, including appendicularians and cnidarians (Luo et al. 2014; Greer et al. 2015; Ohman et al. 2018), colonial bacteria (Davis et al. 1992b) or test-bearing rhizarians (Dennett et al. 2002; Biard et al. 2016; Nakamura et al. 2017; Gaskell et al. in review), which are typically severely disrupted by conventional sampling methods. In addition to unreliably sampling fragile organisms, even vertically stratified plankton nets (e.g., MOCNESS, MULTINET; Wiebe and Benfield 2003) fail to resolve the fine vertical distribution patterns of planktonic taxa. In contrast, towed and autonomous in situ imaging systems have successfully revealed fine-scale distribution patterns of zooplankton communities and the influence of environmental variables (Luo et al. 2014; Greer et al. 2015; Faillettaz et al. 2016; Ohman et al. 2018).

In the present study, we use the in situ imaging Underwater Vision Profiler 5 (UVP5) to identify large test-bearing rhizarians (> 600 μm) (Biard et al. 2016) and characterize their ecological preferences and vertical niche distributions. We use in situ images collected during extensive vertical profiles on four research cruises of *California Current Ecosystem* Long-Term Ecological Research (CCE-LTER) program (2008-2016) (Ohman et al. 2013). CCE-LTER is situated in the southern California Current System, a productive eastern boundary current coastal upwelling ecosystem. Notably, the CCE region is characterized by a high diversity of test-bearing rhizarians (136-200 radiolarian species, mainly sampled from small size fractions) (Kling and Boltovskoy 1995) with periodically high abundances/biovolumes of larger cells (> 600 μm) that can represent a substantial fraction of zooplankton communities (Biard et al. 2016), as well as vertical fluxes of carbon (Stukel et al. 2018) and silica (Biard et al. 2018). Vertical profiling was conducted from the euphotic region down to the dark mesopelagic ocean in order to define the ecological preferences of diverse test-bearing rhizarians. We also employ Generalized Additive Models to model the responses of rhizarian abundance to different environmental variables in relation to of habitat depth and time.

## Methods

### In situ analysis of rhizarian populations

In situ measurements of rhizarian populations were acquired between 2008 and 2016 on four process cruises of the *California Current Ecosystem* Long-Term Ecological Research (CCE-LTER) Program (P0810, P1208, P1408, P1604; the first two digits designate the year and the second two the month). While the different process cruises were directed to specific questions (Ohman 2018), they all shared the same quasi-Lagrangian design, whereby water parcels of interest (labelled as a “Cycle”) were sampled intensively while following the same planktonic population for several days (2-5 days; Landry et al. 2009; Ohman et al. 2013). During these Lagrangian experiments, covering a wide range of hydrographic conditions (e.g., coastal upwelling, mesotrophic waters, offshore oligotrophic waters; details in Stukel et al. 2018), we conducted repeated vertical profiles with a CTD-rosette equipped with the in situ imaging system Underwater Vision Profiler 5 (UVP5). The UVP5 is designed to quantify particle concentration and image large planktonic organisms (> 600 μm) at fine vertical scales (5-20 cm vertical resolution; Picheral et al. 2010). In the present study, abundance data were binned in depth intervals of 5 m, while discrete depth was considered for individual vignettes. A total of 205 vertical UVP5 profiles were retained to estimate integrated abundances, vertical distribution and ecological niche of test-bearing rhizarians. Only 154 of these vertical profiles extended to 500 m (i.e., upper mesopelagic). These vertical profiles generated 22,504 individual rhizarian vignettes (i.e., individual image of particles > 600 μm), after processing the raw images as described in Biard et al. (2016, 2018). All images were manually identified and classified by a single person. Upon request, all individual vignettes, their associated metadata (e.g., area, ESD, etc.) and binned abundances are accessible online at Ecotaxa (http://ecotaxa.obs-vlfr.fr; Picheral 2017).

### Environmental data

To assess the influence of environmental variables on rhizarian distributions, we created a matrix of abiotic variables obtained from 1) CTD downcast data and 2) water-column dissolved inorganic nutrients, sampled at discrete depths. Data, along with detailed sampling and analytical methods, can be found at http://oceaninformatics.ucsd.edu/datazoo/catalogs/ccelter/datasets. From this matrix of environmental data, we used six variables: temperature (° C); dissolved oxygen concentration (μmol kg^−1^); Chl *a* concentration (μg L^−1^); depth of the deep chlorophyll maximum, DCM (m); Chl *a* concentration at the DCM (μg L^−1^); and silicic acid concentration (μmol L^−1^). Temperature was measured using recently calibrated Seabird sensors (SBE 3). Dissolved oxygen was measured using a calibrated SBE 43 polarographic membrane sensor. Chl *a* (http://www.calcofi.org/ccpublications/calcofi-methods/8-chlorophyll-methods.html) and silicic acid concentrations (http://www.calcofi.org/ccpublications/calcofi-methods/422-nutrient-methods.html) were measured on water samples collected at discrete depths with Niskin bottles. Depth of the DCM was determined as the maximum value of in vivo Chl *a* fluorescence detected by the fluorometer attached to the CTD rosette (after a smoothed moving average). The concentration of particles (0.20-27 mm) was recorded by the UVP5. In order to provide silicic acid concentrations at finer vertical resolution than the discrete measurement depths, we performed linear interpolations (0-500 m) for each Cycle.

For each taxonomic group, determined based on common morphological characteristics (see below in Results), we considered the central vertical niche as the median depth (± interquartile range) where vignettes were observed. Subsequently, for these central vertical niches, we computed a mean value for each abiotic variable retained (see above).

### Statistical analysis

All data analysis was conducted using R 3.4.4. (R Core Team, 2018), custom scripts and the following packages: *tidyverse* (Wickham and RStudio 2017) and *visreg* (Breheny and Burchett 2017). We used Generalized Additive Models (GAMs; Wood 2017) implemented in R package *mgcv*, to model the relationship between the abundance of different taxonomic groups of test-bearing rhizarians (log-transformed) environmental variables. As an extension of Generalized Linear Models (GLMs), GAMs offer a powerful and flexible alternative that can estimate linear trends as well as non-monotonic responses. For each taxonomic group, we first performed GAMs with the inclusion of year (i.e., cruise) as predictor to account for interannual (inter-cruise) variations. If no statistically significant effects of cruise were determined, we remove the year as a predictor variable and performed a second GAM. For each model, we obtained the following metrics: fitted values, residuals, adjusted R^2^, restricted maximum likelihood (REML) score, deviance explained (%), and *p* values for F statistics.

## Results

### Diversity of test-bearing rhizarians

Twelve distinct categories were delineated: two types of radiolaria, one foraminiferan, eight phaeodaria and one group of unknown rhizaria (Fig. 1). These categories are hereafter described starting with those inhabiting surface waters, continuing with those dwelling deeper. Collodaria (Fig. 1a) are composed of colonial and single-cell organisms (n = 487 vignettes), and Acantharia (Fig. 1b) displayed their typical star-like shape (n = 931). Neither of these categories was identified to lower taxonomic levels. At the base of the euphotic zone (~100 m), two phaeodarian families were identified: Castanellidae (Fig. 1c) and Aulosphaeridae (Fig. 1d), distinguishable by their spherical morphology. The Castanellidae (n = 814) were characterized by a dark central sphere (malacoma + scleracoma) and small spines at the periphery, the entire central sphere surrounded by an extended network of ectoplasm (grey sphere). On the other hand, the Aulosphaeridae (n = 18,752) possess a geometric meshwork (skeleton) with numerous short spines, and, in its center, a clearly visible phaeodium. A subset of vignettes (n = 188) could not be clearly assigned to one of the 11 specific rhizarian categories and were, in this case, assigned to a general category (Fig. 1e).

**Fig. 1.**
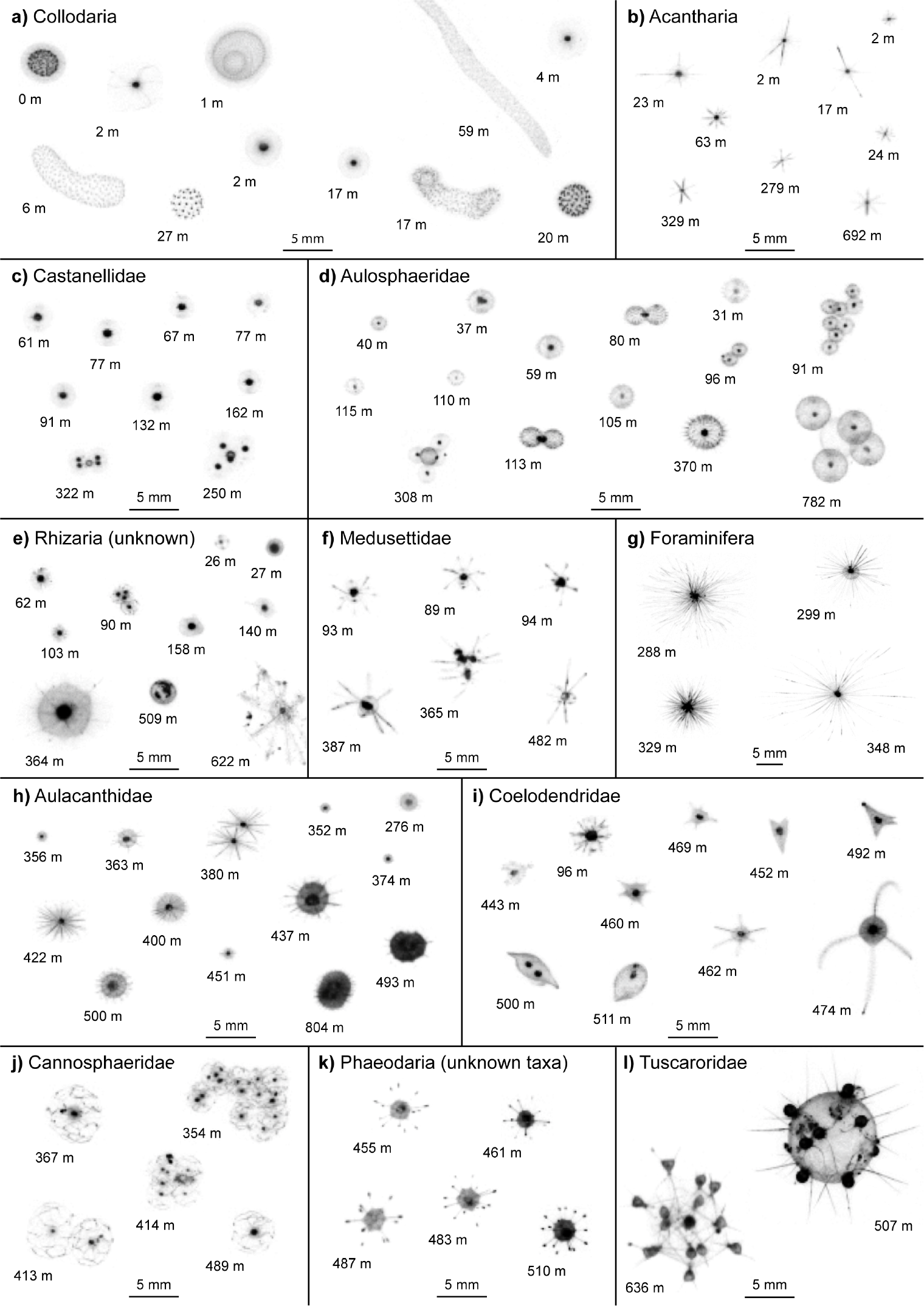
Diversity of representative test-bearing rhizarians obtained with the UVP5 in the *California Current Ecosystem*. Each vignette is displayed with the depth where the organism was observed. All images are scaled the same, except for the Foraminifera.

At greater depths, the phaeodarian family Medusettidae (n = 30) was represented by a few organisms characterized by an ovoid central part from which three pairs of long branch-like extensions (styles) arise (Fig. 1f). Foraminifera (n = 196; Fig. 1g) included unicellular protists with an extended network of pseudopods, reaching up to ten times the length of the central test. The phaeodarian Aulacanthidae (n = 725; Fig. 1h) displayed a continuum of shapes, mainly composed of a spherical/ovoid central part with a phaeodium clearly visible and a variable number of peripherical spines arising from the center. The most diverse phaeodarian family was the Coelodendridae (n = 284), with five unique morphotypes (Fig. 1i), ranging from arrow-like shapes to ovoid cells, all with or without branch-like extensions (styles).

The three remaining categories were restricted to waters deeper than 350 m. First, the Cannosphaeridae (n = 33; Fig. 1j) were characterized by large organisms (2-5 mm) with a loosely constructed geometric meshwork (skeleton) and a small central part in the center. Then, a category of unknown phaeodarians (Fig. 1k) included specimens (n = 61) with an angular cell with 11-14 long spines emerging from the phaeodium. Finally, three specimens were members of the Tuscaroridae (Fig. 1i). These specimens were almost exclusively colonial, with a dozen individual cells displaying 2-3 spicules oriented to the outside, and attached to a large spherical or intricate central structure.

Overall, test-bearing rhizarians detected by UVP5 encompassed organisms ranging in size (expressed as Equivalent Spherical Diameter, ESD) from 877 μm, for a single cell of Castanellidae, to 12.6 mm for a colonial collodarian (Fig. 2). While most categories included organisms with diameters between 1 and 3 mm, several taxa including the Foraminifera (Fig. 1g), Tuscaroridae (Fig. 1i) and Cannosphaeridae (Fig. 1j), mostly exceed a mean size of 3 mm. These three taxa, in addition to colonial collodarians (Fig. 1a), included organisms of large individual sizes (e.g., Cannosphaeridae), with extended pseudopodial webs (e.g., Foraminifera) or forming large colonies (e.g., Tuscaroridae). We observed that not only the preceding taxa, but all phaeodarian families can often be observed as colonies of several unicellular specimens. This colonial feature explains some of the size variability within a single taxon (e.g., Aulosphaeridae).

**Fig. 2.**
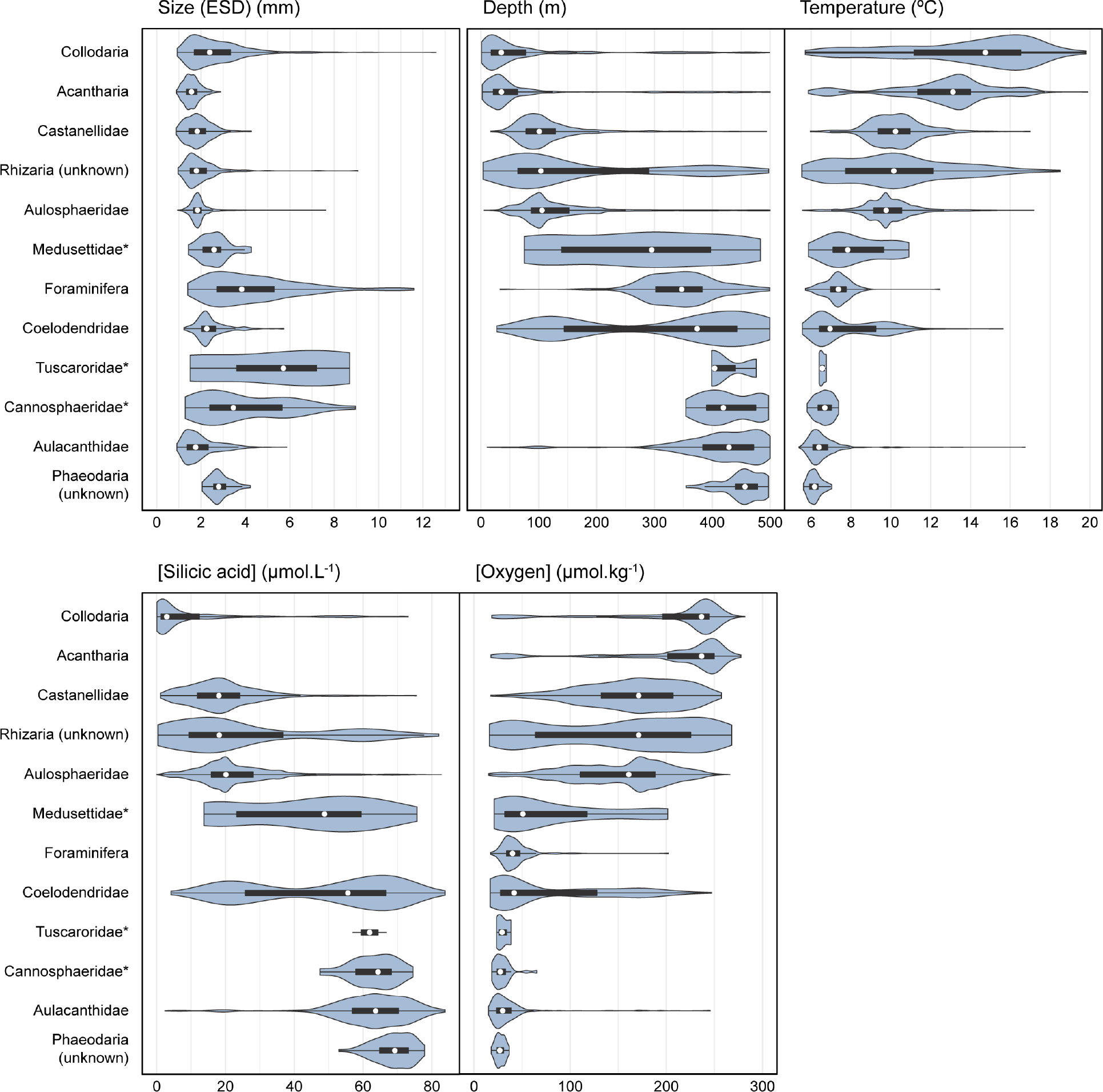
Niche definition of test-bearing rhizarians. Violin plots showing the size distribution and affinity of different rhizarian taxa with four environmental variables. Boxplot indicates median (white dot), ranges between the first and third quartiles, and whiskers cover extreme data points within 1.5 IQR range from the quartiles.

### Vertical niche definition

In order to define test-bearing rhizarians’ realized ecological niches, we compared their vertical distribution with profiles of environmental variables, including temperature, silicic acid and dissolved oxygen (Fig. 2). We defined three layers occupied by specific taxa (Fig. 3a). First, the upper ocean or euphotic zone was characterized by the presence of Collodaria (median vertical niche = 35 m) and Acantharia (35 m), being found primarily in the first 100 m (but sporadically to 500 m). The two taxa occurred at temperatures ranging from > 11° C to 14-16° C, for Acantharia and Collodaria respectively, and thrived in well oxygenated and silicic acid-depleted surface waters. Then, at the base of the euphotic zone and near the upper mesopelagic, we observed three taxa, the Castanellidae (101 m), the unknown Rhizaria (103 m) and finally the Aulosphaeridae (106 m). The three categories were mainly found in well oxygenated (> 100 μmol kg^−1^) water, with temperature records of 10 ± 1° C and silicic acid concentrations of ~20 μmol L^−1^. Finally, the third layer, corresponding to the upper mesopelagic (200-500 m), was characterized by a more diverse community (7 taxa) of test-bearing rhizarians associated with colder temperatures (< 8° C). Two taxa were transitional between the limit of the euphotic zone and the upper mesopelagic: Medusettidae were observed consistently between 150 and 400 m, while the Coelodendridae displayed a bimodal vertical distribution with higher densities at ~100 and ~400 m. Apart from these two specific taxa, the remaining ones (i.e., Foraminifera, Tuscaroridae, Cannosphaeridae, Aulacanthidae and the unknown phaeodarians), were associated with deep (> 300 m) mesopelagic water masses, enriched in silicic acid (> 60 μmol L^−1^) and close to the Oxygen Minimum Zone (OMZ; oxygen concentration < 50 μmol kg^−1^). Overall, we observed an increase of morphotype diversity with depth (Fig. 3b), with substantially larger changes diversity in the epipelagic and the upper mesopelagic.

**Fig. 3.**
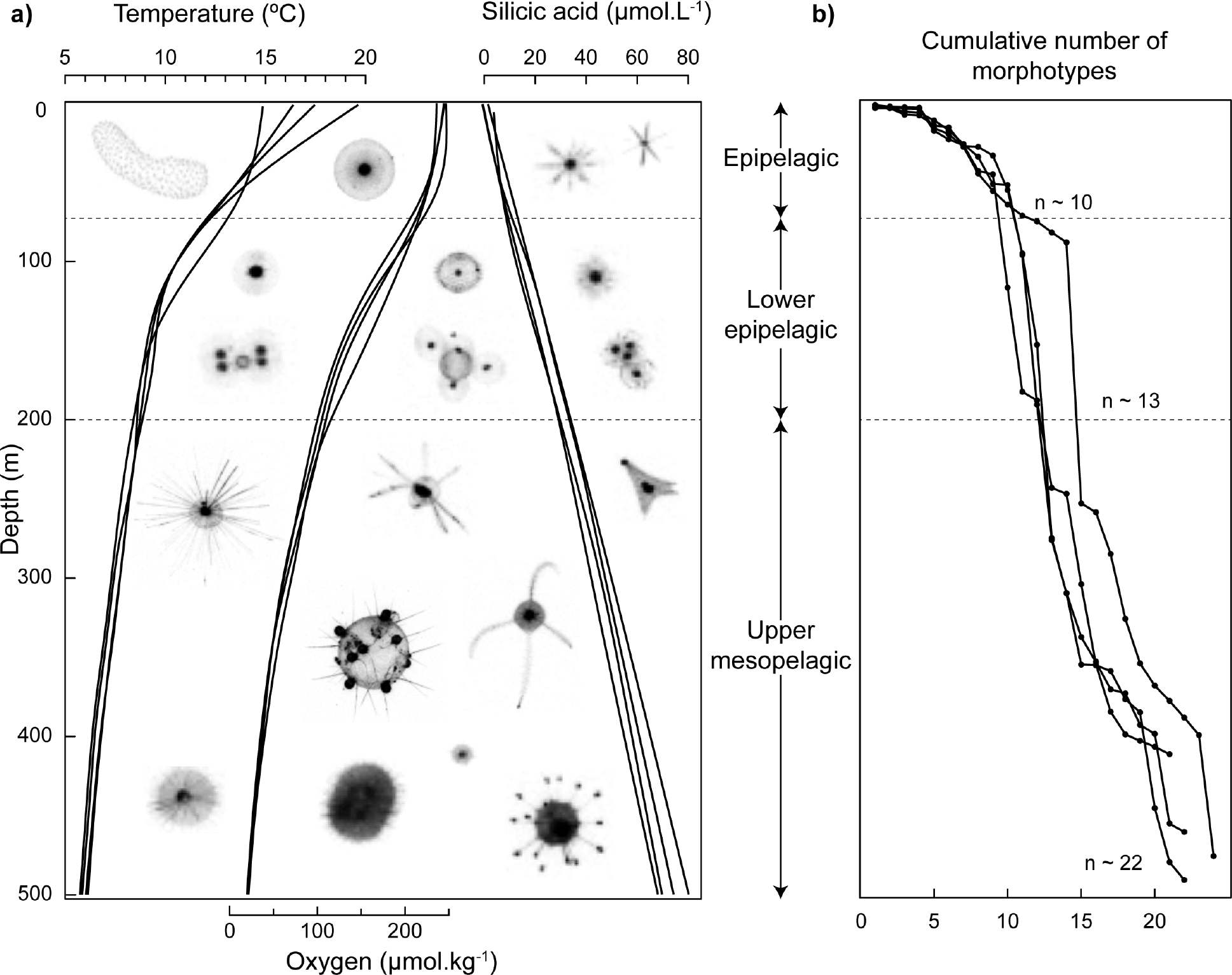
Vertical niche zonation of test-bearing protists (Rhizaria) into the twilight zone of the *California Current Ecosystem*. (**a**) Average vertical profiles (4 lines, one for each cruise) of temperature (left), oxygen concentration (middle) and silicic acid concentration (right) are displayed alongside representative rhizarian taxa. (**b**) Vertical distribution of rhizarian diversity (expressed as number of individual morphotypes). Numbers (n) at the bottom of each zone represent the average cumulative number of morphotypes.

### Rhizaria abundance over time

Integrated abundances of the different rhizarian categories (with the exception of the Tuscaroridae, which were too rare to be reliably sampled) were quantified in the different Cycles and frontal studies (Fig. 4; Supplementary Table ST1). Considering all four cruises, Aulosphaeridae phaeodarians (Fig. 4e) were the most abundant test-bearing rhizarian taxa within the first 300 m (but also extending to 0-500 m). In the upper 300 m, the remaining taxa, Collodaria (Fig. 4a), Acantharia (Fig. 4b) and Castanellidae (Fig. 4c) showed integrated abundances of the same order of magnitude as one another. When extending to the mesopelagic (0-500 m) off California, the abundances of phaeodarians (Fig. 4d-g, i) and foraminiferans (Fig. 4h) were also of the same order of magnitude, with the phaeodarian Aulacanthidae consistently being the most abundant overall.

**Fig. 4.**
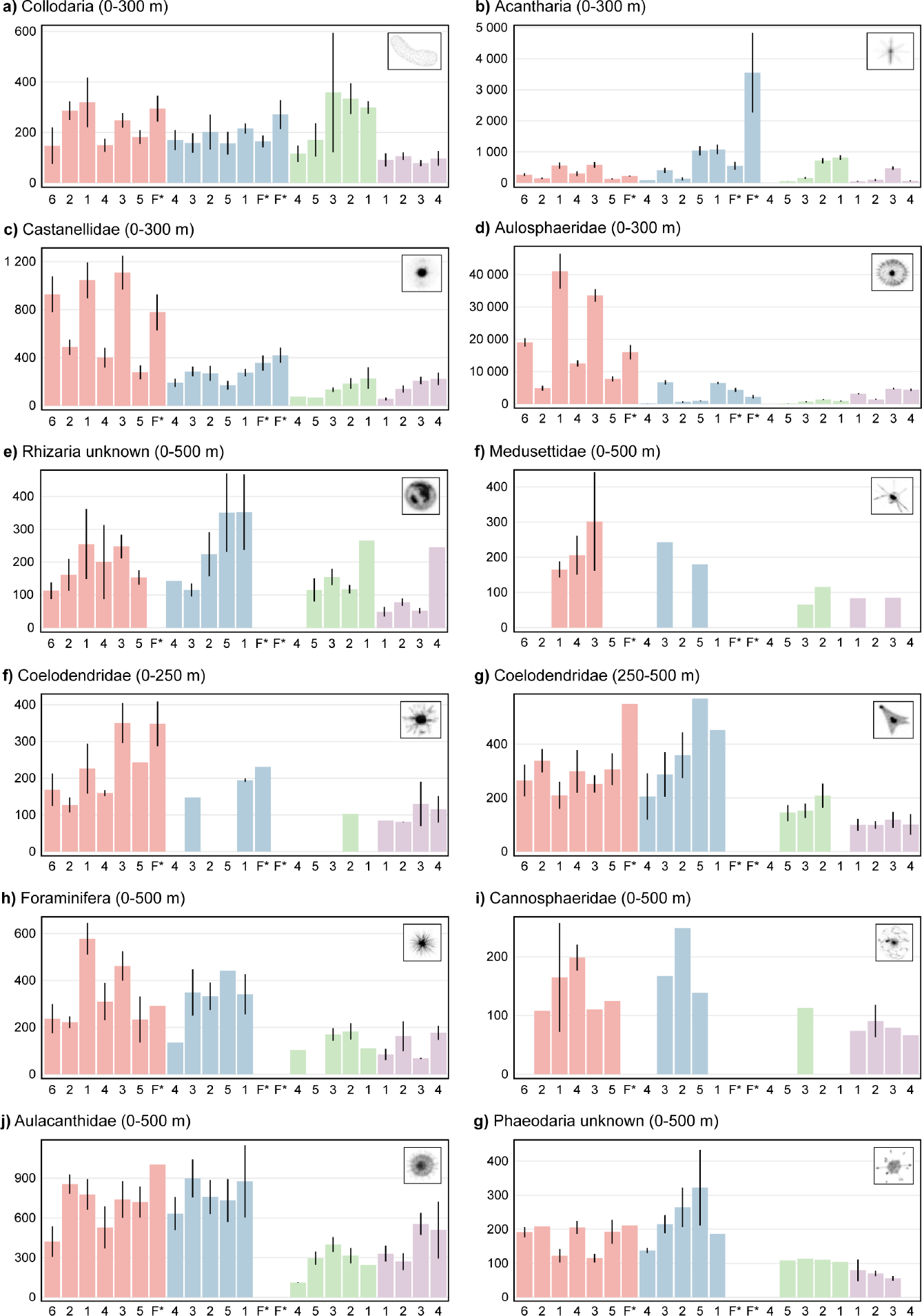
Variation of rhizarian integrated abundances over the four process cruises (red: 2008; blue: 2012; green: 2014; purple: 2016). Depth of integration is displayed in panel headers. Error bars denote standard error of the mean. Within each year, Lagrangian Cycles are ordered from left to right along a gradient of increasing primary production (frontal studies - F- are not considered). Note the independent scaling for each panel. Detailed values in Supplementary Table 1.

The variability over time (i.e., within and between cruises) differed markedly among different types of test-bearing rhizarians and appeared slightly more pronounced in the upper ocean layer. Indeed, for the Castanellidae and the Aulosphaeridae (Fig. 4c, d), abundances were considerably higher and more variable during the 2008 cruise (late summer/early fall) than for the three following cruises (spring and summer). For other taxa primarily occupying upper ocean waters (Collodaria and Acantharia; Fig. 4a, b), inter- and intra-cruise variability appeared reduced with occasionally higher abundances in specific samples. Considering now the full water column down to 500 m, inter-cruise variability appeared less pronounced or negligible for rhizarians living at depth. Although we observed a diminution in integrated abundances between the 2008 + 2012 and the 2014 + 2016 cruises, this difference applied to almost all taxa, including those dwelling near the surface.

### Influence of environmental variables on rhizarian populations

Having established the vertical niches of test-bearing rhizarians and their variability in abundance among cruises, we explored environmental variables that may explain and predict this variability. Generalized Additive Models (GAMs) modeled individual responses of the different taxa (Figs. 5-7) with variable statistical power (0.08 > R^2^_adj_ > 0.92). For each GAM output, we provide a smoothed spline (reduced to a simple linear effect if necessary) of each variable’s relative contribution to predicting observed abundances.

**Fig. 5.**
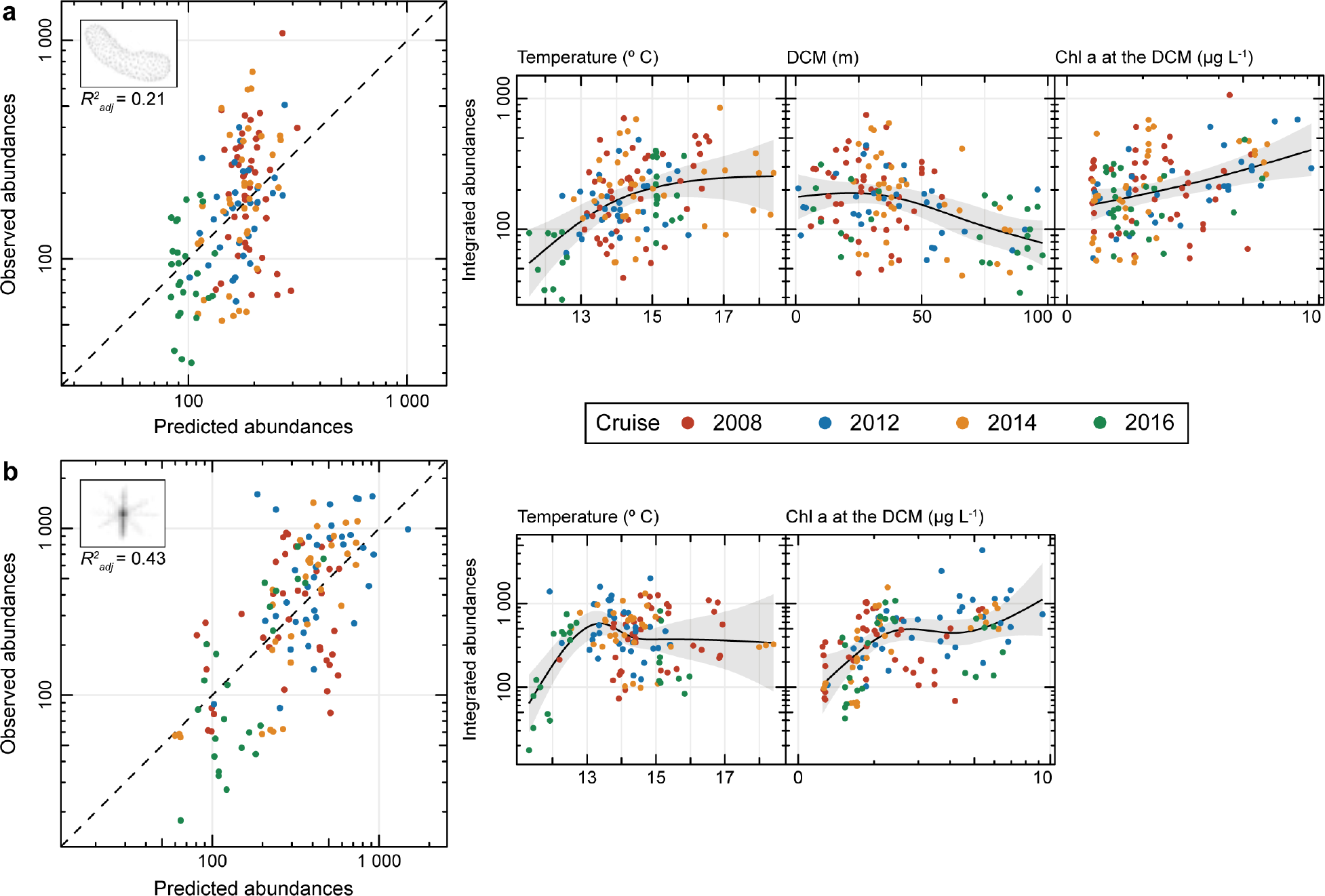
Generalized Additive Models (GAMs) based on selected niche abiotic characteristics of epipelagic rhizarians: **(a)** Collodaria and **(b)** Acantharia. Grey intervals denote the 95% confidence intervals for each spline. Adjusted R-squared for the models are indicated for each taxon.

For the epipelagic zone, temperature, the depth of the Deep Chl *a* Maximum (DCM) and Chl *a* concentration [Chl *a*] at the DCM were retained as explanatory variables for the GAMs, and produced R^2^_adj_ of 0.43 and 0.21, for Acantharia and Collodaria, respectively. No significant effect of sampling year was recorded for either taxon. The predictive power for the Collodaria GAM model was low (R^2^_adj_ = 0.21), but all three tested variables had significant effects (Fig. 5a). For Acantharia (Fig. 5b), the DCM appeared to have an insignificant effect in the model, while temperature was the most significant variable. Both taxa showed the same response of increasing abundances with increasing [Chl *a*] at the DCM. The response to increasing temperatures revealed a positive and marked response from 11°C to 13-14° C with increasing abundances of both taxa, followed by a reduced or negligible response (i.e., threshold effect). Only the Collodaria displayed a significant negative response to the deepening of the chlorophyll maximum (Fig. 5a).

For rhizarians dwelling at the basis of the epipelagic zone, temperature, DCM, particle concentration and silicic acid were explanatory variables. Carbon fluxes (recorded with sediment traps at 100 m) show no significant effects on abundances of both types of organisms (Castanellidae, τ = 0.03; *p* = 0.60; Aulosphaeridae, τ = 0.07; *p* = 0.25) and was not considered further. The resulting GAMs produced moderate to strong R^2^_adj_ values of 0.71 and 0.92, for Castanellidae (Fig. 6a) and Aulosphaeridae (Fig. 6b), respectively. Here, year/cruise had a significant and strong effect on observed abundances. The contribution of each variable was highly variable and produced inconsistent patterns for individual taxa, with the year 2014 standing out with often opposite patterns. The resulting models suggested: a minimal effect of the DCM on both taxa; a limited effect of increasing particle concentration, except for the sharp increase of abundance of Aulosphaeridae in 2012; a general decrease in abundance with increasing temperature (excepted for 2014). The modelled response to increasing silicic acid concentration differed between Castanellidae and Aulosphaeridae, with the former showing a negative or negligible response, while the later showed a weakly positive response.

**Fig. 6.**
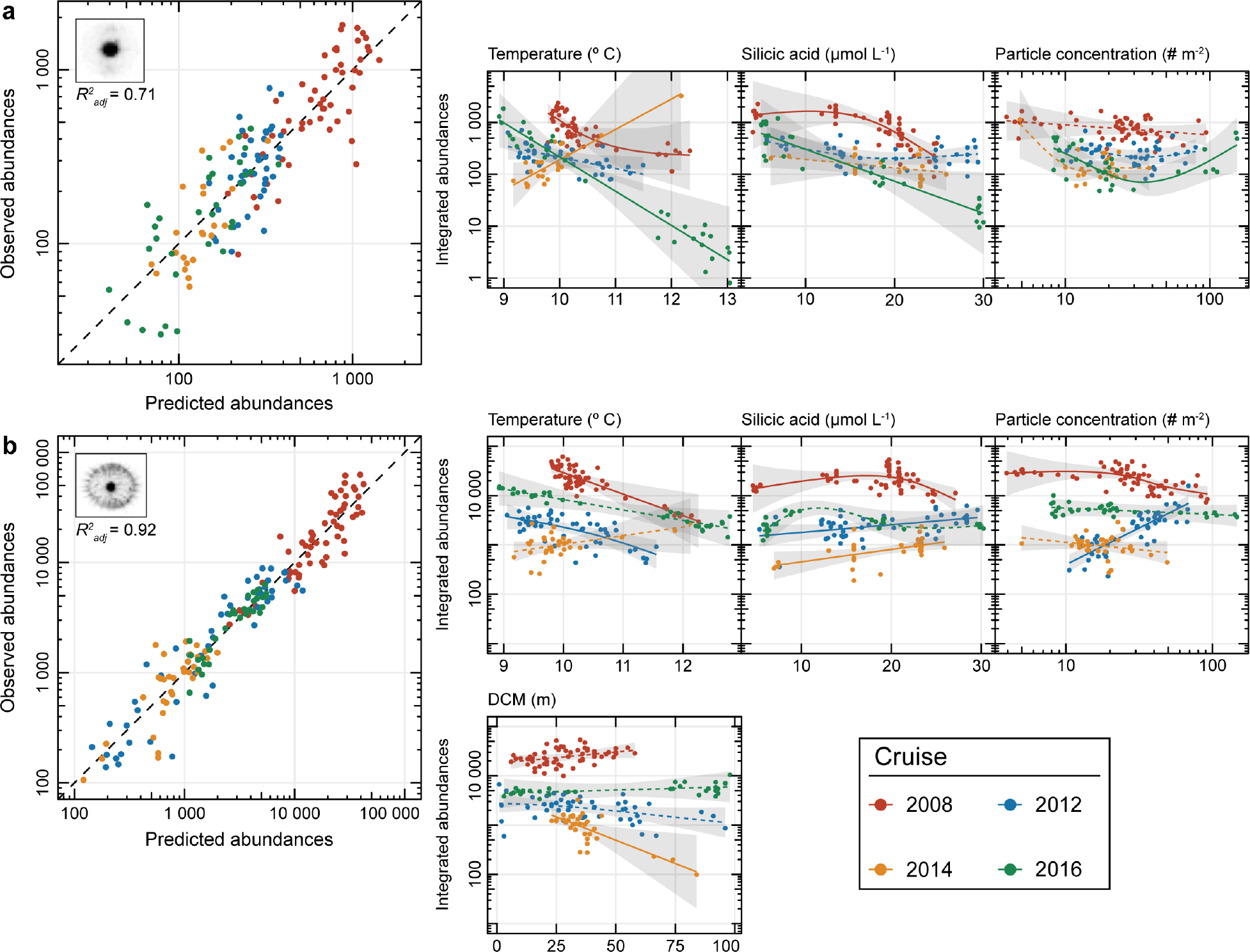
GAMs based on selected niche abiotic characteristics of lower epipelagic rhizarians: (**a**) Castanellidae and (**b**) Aulosphaeridae. For sub panels, solid lines represent significant GAMs and dashed lines, insignificant GAMs. Grey intervals denote the 95% confidence intervals for each spline. Adjusted R-squared for the models are indicated for each taxon.

In the third zone, i.e., the upper mesopelagic layer, GAMs produced models with weak-to-moderate predictive power: R^2^_adj_ < 0.3 for Foraminifera, Aulacanthidae and Coelodendridae; R^2^_adj_ = 0.51 for the unknown phaeodarians (Fig. 7). Similar to the epipelagic GAMs, we detected no significant effects of sampling year on observed abundances of these 4 taxa, nor considered Carbon flux at 100 m as explanatory variable. The only significant, albeit weak, response of Foraminifera to an abiotic factor was a decrease in observed abundance with increasing dissolved oxygen (Fig. 7b). The three remaining taxa for this zone showed consistent responses, starting with the Coelodendridae near the surface and their threshold response to temperature (similar to Acantharia and Collodaria; Fig. 5). For the phaeodarians thriving in the upper mesopelagic (Fig. 7a, c, d), abundances consistently increased with increasing silicic acid concentrations.

**Fig. 7.**
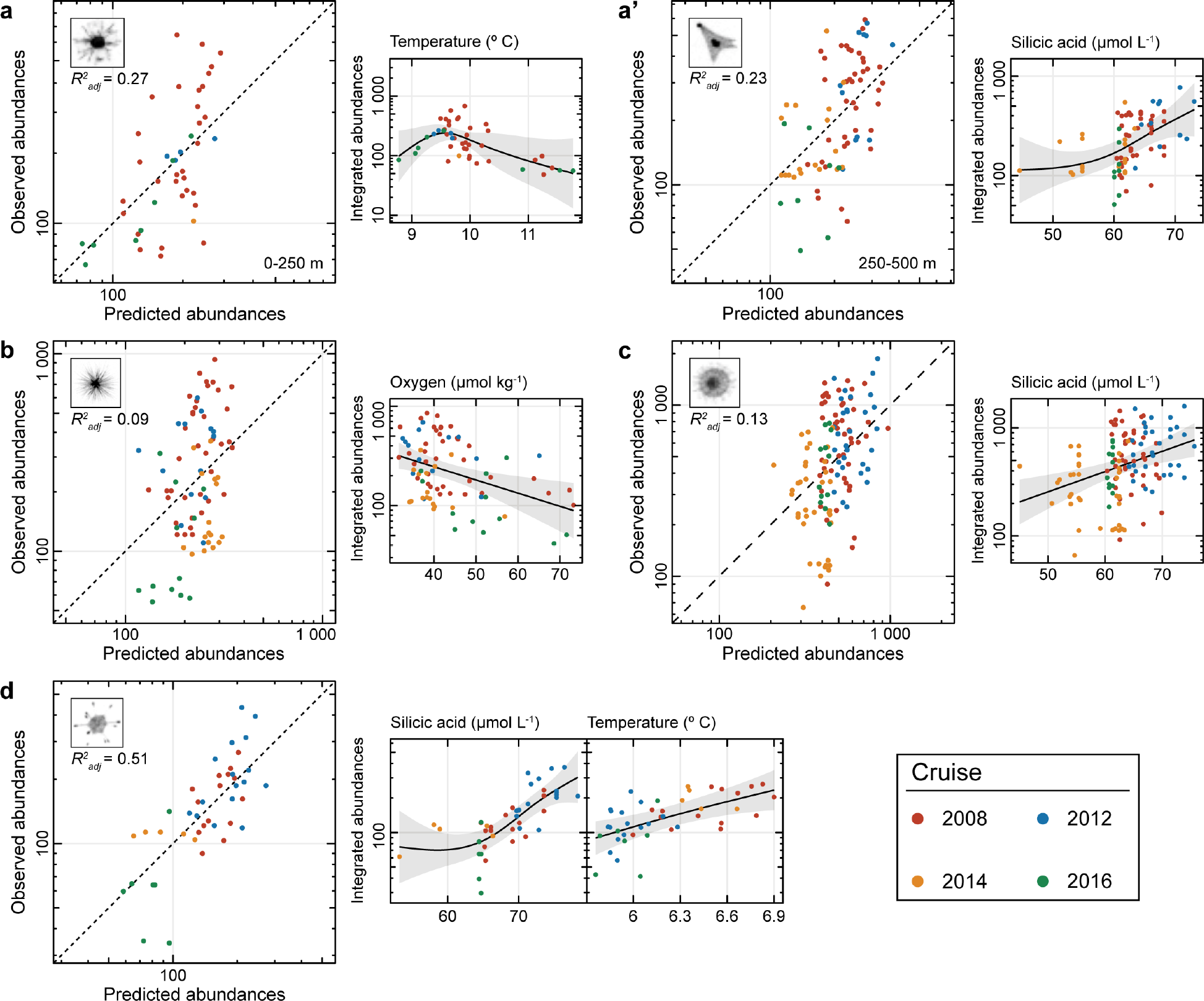
GAMs based on selected niche characteristics of mesopelagic rhizarians: (**a-a’**) Coelodendridae for a depth range of 0-250 m (left panels) and 250-500 m (right panels), (**b**) Foraminifera, (**c**) Aulacanthidae and (**d**) Phaeodaria unknown. Grey intervals denote the 95% confidence intervals for each spline. Adjusted R-squared for the models are indicated for each taxon.

## Discussion

### Diversity of test-bearing rhizarians from in situ imaging

In situ images of test-bearing rhizarians in the *California Current Ecosystem* revealed 12 visually-recognizable morphological categories, with mixed taxonomic ranks inherent to the minimum size resolvable by the UVP5 (> 600 μm). Early studies in the CCE and nearby regions reported radiolarian diversity an order of magnitude higher, ranging from 136 to 200 species (Casey 1966; Boltovskoy and Riedel 1987; Kling and Boltovskoy 1995). These estimates of diversity cannot be directly compared with the present results, as early studies used plankton nets (with mesh size ≤ 62 μm) targeting the small polycystines that are undetectable with the UVP5, and also analyzed preserved specimens by microscopy. However, since rhizarian tests can be easily damaged upon collection with plankton nets (Nakamura and Suzuki 2015; Suzuki and Not 2015) or dissolved upon preservation with traditional fixatives (Beers and Stewart 1970), in situ imaging can confer other advantages. In situ imaging is a non-invasive means of observing fragile planktonic organisms (e.g., Remsen et al. 2004; Benfield et al. 2007), including test-bearing rhizarians, in their natural habitat. Such images can record natural feeding postures and extension of rhizopodia, the organisms’ 3-dimensional orientation, aggregations of multiple cells, etc. (Dennett et al. 2002; Biard et al. 2016; Ohman et al. 2018; Gaskell et al. in review). For non-rhizarian taxa, imaging has resolved interactions between predators and prey (Ohman et al. 2018) and fine-scale distributions (Luo et al. 2014; Faillettaz et al. 2016).

While seven of the 12 rhizarian categories observed here were previously described in this region, others, including the phaeodarians Medusettidae, Coelodendridae, Cannosphaeridae and Tuscaroridae have rarely been reported in the CCE, and in few instances worldwide. Among these taxa, Cannosphaeridae are often captured in subtropical waters but are severely damaged upon collection (Nakamura and Suzuki 2015). We could not find any reference to the Medusettidae in contemporary studies off California and only a few worldwide (e.g., Cachon and Cachon-Enjumet 1965). Only a handful of studies have reported the presence of large colonial Tuscacoridae (Ling and Haddock 1997) and Coelodendridae (Zasko and Rusanov 2005). For these taxa, in situ collection with Remotely Operated Vehicles (Beittenmiller 2015), submersible vehicles (Swanberg et al. 1986) or in situ imaging (Nakamura et al. 2017) has enabled detailed observations. Regardless of differences in taxonomic resolution between this and previous studies, our detection of these elusive taxa at depth supports the increase in rhizarian diversity with depth near the CCE (Gulf of California; Zasko and Rusanov 2005) or elsewhere (e.g., highest phaeodarian diversity at ~7000 m in the Kamchatka, Reshetnyak 1955). This increase in diversity is inconsistent with the expected decrease in protistan diversity toward the mesopelagic (Robinson et al. 2010) but should be viewed as an example of adaption of phaeodarians to deep-dwelling life. We have also confirmed the tendency for larger organisms to occur deeper in the water column, a typical feature not only of phaeodarians (Zasko and Rusanov 2005), but also numerous other deep-sea organisms (e.g., Bergmann’s principle of increasing cell size with decreasing temperature; Timofeev 2001).

### Vertical niche distribution and influence of environmental factors

With images taken in situ every ~5-20 cm and binned to depths of 5 m, from the surface down to the mesopelagic layer, we were able to define the vertical niches of different taxa of test-bearing rhizarians at unprecedented resolution. Until the advent of in situ imaging techniques, vertical niche definition relied on the use of traditional sampling methods like plankton nets, Niskin bottles or pumps, all representing rather coarse vertical resolution, leading to broad estimates of vertical niches (Dworetzky and Morley 1987; Zasko and Rusanov 2005). Although four dedicated studies have used advanced imaging technologies for rhizarian habitat studies (Dennett et al. 2002; Biard et al. 2016; Nakamura et al. 2017; Gaskell et al. in review), only Gaskell et al. (in review) resolved fine-scale vertical distributions and relations to environmental variables for test-bearing rhizarians, besides the present study. Overall, our results are surprisingly similar to previous assessments of the vertical ranges of polycystine radiolarians in the northern subtropical Pacific (Boltovskoy et al. 2017), although most of the organisms detected in the present study belong to the Phaeodaria. In Boltovsky et al. (2017), four major vertical zones were delineated: (1) species limited to surface waters (< 100 m), (2) species inhabiting the base of the euphotic zone (~100 m), (3) species from intermediate waters (100-300 m) and (4) deep-dwelling species (> 300 m). We discuss vertical zonation further below.

### Epipelagic zone

In the epipelagic layer (i.e., 0-100 m), both Collodaria and Acantharia were the dominant taxa, along with a few other less common forms. The presence of both of these taxa in surface layers have been reported on several occasions and in contrasting ecosystems (e.g., Beers and Stewart 1971; Anderson 1983; Michaels 1988; Boltovskoy et al. 2017). This affinity with surface illuminated layers is likely related to the presence of photosymbionts in some acantharian cells (i.e., in 3 out of 9 molecular clades; Decelle and Not 2015) and in almost all collodarian species (Hollande and Enjumet 1953). Notably, the nonsymbiotic relatives of acantharians (Decelle and Not 2015) and collodarians (probably limited to the family Collophidiidae; Biard et al. 2017) have been observed deeper in the dark ocean (Schewiakoff 1926; Bernstein et al. 1999; Biard et al. 2017). Interestingly, although both have the ability to adjust their buoyancy and vertical position (Michaels 1988), physical mechanisms at the air-water interface (e.g., Langmuir cells or *zoöcurrent* as called by Haeckel 1893) can lead to increased concentration or patchiness in surface layers (Casey 1971; Swanberg 1983; Michaels 1988). This phenomenon could explain high acantharian densities observed in frontal regions in 2012 (Fig. 4b), a similar pattern observed in situ in a frontal zone of the Ligurian Current (Mediterranean Sea; Faillettaz et al. 2016).

In the surface layer, temperature appeared as the main environmental factor related to the abundance of collodarians and acantharians, with lower abundances in waters cooler than 13-15° C. This pattern is consistent with previous studies, highlighting the role of temperature (followed by nutrients and primary productivity) in this layer (Dworetzky and Morley 1987; Boltovskoy and Correa 2016). While increases in collodarian abundances have been previously associated with increasing chlorophyll *a* concentration at the deep chlorophyll maximum (Faillettaz et al. 2016), here a deeper DCM also led to a decrease in abundances. Whether this is a cause-and-effect relationship is unclear, as chlorophyll *a* concentration at the DCM was negatively correlated with the depth of the DCM (τ = −0.38; *p* < 0.05). Further analyses should evaluate the potential of these two environmental variables as predictors for the abundance of upper ocean rhizarians.

The phaeodarians Aulacanthidae were, surprisingly, not commonly observed in surface layers, while they can represent one of the most common phaeodarian groups found in the euphotic layer (e.g., genus *Aulacantha*; Cachon-Enjumet 1961). Instead, here we observed them deeper, in the upper mesopelagic (discussed further below). Although we cannot rule out the existence of two distinct populations associated with either euphotic or mesopelagic layers, their putative absence here might be a consequence of the limited resolution of the UVP5, since most *Aulacantha* are smaller than our detection limit of 600 μm.

### Lower epipelagic zone

At the base of the epipelagic zone, the CCE region was characterized by two abundant phaeodarian taxa, Aulosphaeridae and Castanellidae, and other less abundant rhizarians, including the Coelodendridae. Further south, in the Gulf of California, their presence has been previously observed at similar depths, expected for the Aulosphaeridae, which were observed deeper and with considerably lower abundances (Zasko and Rusanov 2005). Elsewhere, these taxa have also been observed in slightly deeper layers (e.g., < 500 m in the South Atlantic; Kling and Boltovskoy 1995) where they can occupy a significant fraction of zooplankton biomass (e.g., 2.7-13% at 200-300 m at station ALOHA; Steinberg et al. 2008). However, it has been hypothesized that the vertical distribution of test-bearing Rhizaria can vary with latitude, with some species found in shallow surface waters in high latitudes, but potentially in deeper layers in low latitudes (Casey 1977). This variability is believed to be related to water temperature, a factor constraining phaeodarian vertical distributions elsewhere (reviewed in Nakamura and Suzuki 2015). In a companion study to the present work, Aulosphaeridae were inversely correlated with the depth of the 10° C isotherm (Stukel et al. 2018). Here, we generally observed a significant decrease of lower epipelagic phaeodarian populations (Aulosphaeridae and also Castanellidae) with increasing temperature at the depth of their preferred habitat (~100 m). In addition to water temperature, food supply and silicic acid concentration are secondary limiting factors for phaeodarians (reviewed in Nakamura and Suzuki 2015).

Early on, it was hypothesized that deep-water rhizarians should be more abundant below regions of high productivity (Casey 1987). We did not find strong evidence of increased phaeodarian abundance with particle concentration (considered here as a proxy for the food of flux-feeders). Neither were relationships detected between the abundance of phaeodarians and the magnitude of carbon export, as determined using sediment traps or the ^234^Th method. However, phaeodarians, known to be flux feeders (Gowing 1989), are also omnivores that feed on suspended bacteria and protists (Anderson 1983; Gowing 1986; Gowing and Wishner 1992). As little is known about phaeodarian ecology (Nakamura and Suzuki 2015), the relative importance of these two different diets is unknown. We cannot exclude the possibility that a lack of response to increased particle concentrations reflects a predominantly omnivorous diet in the lower epipelagic.

The two phaeodarian taxa inhabiting the lower epipelagic showed contrasting responses to increased silicic acid concentrations, from a slight decrease in abundance of Castanellidae to moderate increases for the Aulosphaeridae. For smaller phaeodarians, the availability of silicic acid (i.e., dSi) could influence their vertical distribution, since some species migrate to layers of higher silicic acid concentrations (Okazaki et al. 2004). Here we did not find evidence of changes in vertical distribution over time, suggesting a rather stable positioning at ~100 m (see also Ohman et al. 2012), potentially corresponding to a habitat that optimizes phaeodarian temperature preferences and food availability. In this scenario, silicic acid concentration would have a limited effect on lower epipelagic rhizarian communities, contrasting with observations from the upper mesopelagic.

### Upper mesopelagic zone

Below the 200 m horizon in the CCE, in situ observations revealed a diverse rhizarian community, characterized mostly by large phaeodarians and planktonic foraminiferans, both with relatively stable abundances over time. Unlike their relatives from the lower epipelagic zone, the three testate phaeodarian taxa, Coelodendridae, Aulacanthidae and unknown phaeodarians, displayed a significant increase of abundance with increasing dSi in the mesopelagic layer. Together with the lack of or weak response to temperature and particle concentrations, this pattern is the first direct evidence that dSi availability may be the most important environmental variable (among those tested) structuring deep phaeodarian communities. Below a certain depth threshold, the decrease in water temperature (shown early on to be a major variable affecting phaeodarian vertical distribution; Nakamura et al. 2013) is slow and constant. Therefore, below a given temperature, phaeodarians are likely to meet favorable conditions to survive and grow at depth, but our results suggest that their success will then be constrained by dSi availability.

In this layer, in situ observations also revealed that various test-bearing rhizarians thrive in oxygen depleted water (< 50 μmol kg^−1^). Only digitate foraminifera (e.g., *Hastigerina* spp.), observed in the present study, have already been associated with oxygen minimum zone (OMZ; Hull et al. 2011; Gaskell et al. in review), while a similar pattern has not been reported so far for phaeodarians. This association with the OMZ was believed to coincide with a peak in mesopelagic biomass, located above the OMZ, upon which the carnivorous digitate foraminifera depend for locating their prey (e.g., calanoid copepods; Hull et al. 2011). Given their omnivorous behavior (Anderson 1983; Gowing 1986; Gowing and Wishner 1992), in situ observations of specimens covered with marine snow (Hull et al. 2011), and the lack of in situ images that could support hypothesis of feeding by phaeodarians on metazoans, it seems likely that phaeodarians are less dependent on the OMZ position than their carnivorous relatives.

### Insights into rhizarian ecology and evolution

In the upper layers of the CCE (0-200 m) most test-bearing rhizarians showed pronounced changes in abundances within and between cruises. Early on, it was reported that the upper 200 meters off Southern California showed seasonal changes in rhizarian populations, depending on ocean circulation (Casey 1966). Notably, such seasonal fluctuations in rhizarian assemblages were not reported deeper, in the mesopelagic layer (Casey 1971), a consistent pattern observed here. Further analyses should focus on resolving depth-dependent rhizarian population variability in the CCE with respect to ocean circulation and upwelling dynamics (e.g., Lavaniegos and Ohman 2007).

Although we successfully modelled the effects of several environmental variables using GAMs, we cannot rule in or out the impact and importance of biotic interactions in explaining distribution patterns and seasonal variations. Little is known about potential predators, parasites, or pathogens of test-bearing rhizarians. The few predators reported so far include hyperiid amphipods (Swanberg and Harbison 1979) or lobster larvae (O’Rorke et al. 2012), while other taxa have been shown to avoid the consumption of rhizarians, potentially because of sterol compounds produced by photosymbiotic species (Swanberg 1979; Anderson 1983). To our knowledge, no comprehensive reports of test-bearing rhizarians have been made from the gut contents of mesopelagic metazoans. Therefore, although our models explained a moderate percentage of deviance, our limited knowledge of biotic interactions with test-bearing rhizarians currently prevent the inclusion of biotic drivers in models.

The present data provide insights into the evolutionary processes of test-bearing rhizarians with respect to their environment and competition with other silicifiers, diatoms in particular. Although radiolarians were among the first protistan lineages inhabiting the primitive ocean, the decrease in dSi concentration during the Cenozoic (ca. 66 Ma), coupled with the rise of diatoms, led to a decrease in radiolarian silicification and ecological success (Lazarus et al. 2009). Here we distinguished three distinct vertical layers, that may reflect different biological adaptations. In surface layers of the CCE, where diatoms often dominate phytoplankton communities (e.g., Brzezinski et al. 2015; Taylor et al. 2015), two test-bearing rhizarians dominate in dSi poor water masses: 1) Collodaria, the most recent radiolarian lineage (ca. 40 Ma; Suzuki and Oba 2015), with most species lacking silicified structure (Biard et al. 2015) and 2) Acantharia, which build tests of strontium sulfate. Both forms are therefore not influenced (or minimally influenced for the few silicified collodarians) by the lack of dSi. Deeper, from the base of the euphotic zone to the deep and dark oceanic layers where diatoms are absent, dSi concentrations increase. There, large silicified phaeodarians become more diverse with depth and their abundance is likely influenced by dSi availability. These results suggest the importance of silica in structuring planktonic communities from the surface to the deep ocean.

## Conclusions

Interest in planktonic rhizarians has expanded markedly in the last two decades, thanks to the development of molecular techniques and in situ imaging (Caron 2016). Using non-invasive in situ imaging, here we found three primary vertical habitats of test-bearing planktonic Rhizaria in the upper 500 m of the water column off Southern California: epipelagic, lower epipelagic, and upper mesopelagic assemblages. The taxonomic richness of planktonic Rhizaria increases in the mesopelagic zone, where large silicified specimens are prevalent. The abundance of the dominant rhizarians in each of these zones is largely defined by: **Epipelagic**-temperature and Chl-*a* at the DCM; **Lower epipelagic**-temperature and silicic acid concentrations; **Upper mesopelagic silicifiers**-silicic acid; **Upper mesopelagic Foraminifera**-dissolved oxygen. Future studies should also treat the role of biotic interactions.

While the present study identifies the habitats of different categories of rhizarians, companion studies in this region evaluated some of their biogeochemical impacts. Large phaeodarian Aulosphaeridae, together with Castanellidae, transport important amounts of biogenic silica to the deep ocean (Biard et al. 2018) and also play substantial roles in carbon flux attenuation in the mesopelagic (Stukel et al. 2018). Together, these studies suggest the need to represent larger planktonic Rhizaria in ecological and biogeochemical models, with explicit consideration of their depth-dependent impacts.

## Acknowledgments

This work was supported by the Scripps Postdoctoral Fellowship program and by National Science Foundation grants OCE-1637632 and OCE-1614359 to the CCE-LTER site. We thank to Y. Nakamura and N. Suzuki for their help with the taxonomic outline of Rhizaria. We are grateful to M. Picheral for his help handling the UVP5 data and the operators who conducted UVP5 deployments during CCE-LTER process cruises. This study is also a contribution from the CHAIRE Vision, supported by the CNRS and Sorbonne University and the French Investissements d’Avenir program.

