## Supplemental Table 1 for "Vertical niche definition of test-bearing protists (Rhizaria) into the twilight zone revealed by in situ imaging"

**Supplementary Table 1.** Integrated abundances (mean ± standard error of the mean) of test-bearing rhizarians in four different cruises.

| **Year** | **Site** | **Collodaria**  **(0-300m)** | **Acantharia**  **(0-300m)** | **Castanellidae**  **(0-300m)** | **Aulosphaeridae**  **(0-300m)** | **Rhizaria**  **(unknown)**  **(0-500m)** | **Medusettidae**  **(0-500m)** | **Coelodendridae (0-250m)** | **Coelodendridae (0-250m)** | **Foraminifera**  **(0-500m)** | **Cannosphaeridae**  **(0-500m)** | **Aulacanthidae**  **(0-500m)** | **Phaeodaria**  **(unknown)** |
| --- | --- | --- | --- | --- | --- | --- | --- | --- | --- | --- | --- | --- | --- |
| 2008 | C1 | 319±99 | 560±100 | 1044±150 | 41028±5387 | 255±107 | 165±23 | 226±68 | 210±50 | 578±67 | 165±92 | 778±116 | 123±19 |
|  | C2 | 286±37 | 148±27 | 487±149 | 4860±719 | 161±49 | - | 127±20 | 338±43 | 222±25 | 108 | 854±73 | 298 |
|  | C3 | 248±29 | 585±89 | 1108±142 | 33515±1929 | 248±36 | 302±140 | 350±55 | 252±32 | 462±63 | 110 | 739±137 | 116±12 |
|  | C4 | 149±26 | 297±79 | 401±82 | 12479±956 | 201±113 | 206±55 | 160±8 | 299±80 | 310±80 | 199±22 | 528±159 | 206±19 |
|  | C5 | 182±27 | 129±23 | 277±58 | 7719±775 | 153±22 | - | 243 | 306±59 | 233±98 | 124 | 720±117 | 193±35 |
|  | C6 | 147±72 | 264±43 | 926±149 | 19040±1293 | 113±25 | - | 169±44 | 265±59 | 237±63 | - | 421±114 | 192±15 |
|  | F1 | 294±51 | 219±14 | 777±150 | 16023±2255 | - | - | 348±61 | 549 | 292 | - | 1003 | 211 |
| 2012 | C1 | 215±20 | 1075±154 | 274±31 | 6474±336 | 353±115 | - | 194±4 | 452 | 341±86 | - | 876±271 | 187 |
|  | C2 | 201±69 | 136±52 | 269±62 | 624±217 | 224±67 | - | - | 359±85 | 333±59 | 249 | 759±127 | 264±57 |
|  | C3 | 158±38 | 403±85 | 284±40 | 6662±646 | 115±20 | 243 | 147 | 287±83 | 349±99 | 167 | 898±144 | 215±26 |
|  | C4 | 170±40 | 88 | 190±33 | 169±13 | 143 | - | - | 205±87 | 135 | - | 632±125 | 138±7 |
|  | C5 | 157±45 | 1038±149 | 169±37 | 952±142 | 351±120 | 180 | - | 570 | 441 | 139 | 732±162 | 322±111 |
|  | F1 | 164±24 | 551±129 | 355±63 | 4334±632 |  | - | 231 | - | - | - | - | - |
|  | F2 | 271±57 | 3551±1286 | 419±63 | 2216±566 |  | - | - | - | - | - | - | - |
| 2014 | C1 | 299±25 | 816±78 | 226±90 | 388±139 | 266 | - | - | - | 111 | - | 244 | 104 |
|  | C2 | 334±61 | 714±88 | 184±45 | 1348±94 | 117±13 | 116±0 | 102 | 209±45 | 183±35 | - | 315±57 | 111 |
|  | C3 | 358±237 | 162±29 | 132±19 | 706±138 | 155±25 | 65 | - | 152±26 | 170±27 | 113 | 399±57 | 113±0 |
|  | C4 | 116±33 | - | 76 | - | - | - | - | - | 104 | - | 113±2 | - |
|  | C5 | 171±66 | 57±1 | 67 | 166±35 | 115±35 | - | - | 146±30 | - | - | 296±50 | 109 |
| 2016 | C1 | 91±26 | 51±12 | 55±11 | 3180±211 | 49±15 | 83 | 84 | 100±22 | 85±23 | 74 | 330±60 | 80±32 |
|  | C2 | 106±15 | 95±30 | 138±29 | 1349±112 | 78±11 | - | 81±1 | 100±14 | 163±64 | 91±27 | 271±64 | 70±8 |
|  | C3 | 78±12 | 474±54 | 208±30 | 4626±274 | 52±8 | 84 | 130±60 | 119±29 | 68±3 | 79 | 555±82 | 56±7 |
|  | C4 | 97±19 | 68±15 | 222±52 | 4355±359 | 245 | - | 115±36 | 101±38 | 177±30 | 66 | 508±215 | - |
